# GenoME: a MoE-based generative model for individualized, multimodal prediction and perturbation of genomic profiles

**DOI:** 10.64898/2025.12.28.696482

**Authors:** Jiachen Wei, Yue Xue, Hao Chai, Yi Qin Gao

## Abstract

The non-coding genome operates through a complex, multiscale regulatory system where regulated gene expressions are closely associated with cell-type-specific histone modifications, transcription factor binding and 3D conformation. Developing computational models that can integrate these patterns to predict and interpret the regulatory system remains challenging. Here, we present GenoME, a Mixture of Experts (MoE)-based generative model that uses DNA sequence and cell-type-specific ATAC-seq signals to predict a unified genomic profile encompassing epigenomics, transcriptomics, and chromatin architecture at base-pair to kilobase resolutions. GenoME enables multiscale predictions for held-out genomic regions and, critically, generalizes to predict the full regulatory landscape of unseen or individualized cell types from a single ATAC-seq input. We equip GenoME with an *in silico* perturbation framework that accurately forecasts the multimodal consequences of genetic perturbations and identifies functional enhancer-promoter connections, outperforming specialized models like Activity-by-Contact. These predictions can also be used to decipher the transcription factor grammar of cell-type-specific enhancers. GenoME thus provides a versatile, all-in-one platform for generative modeling, cross-cell-type generalization, and causal mechanistic investigation of the multiscale regulatory genome.

## Introduction

The eukaryotic genome is regulated by a dynamic, interconnected system operating across multiple scales^1–5^. The cis-regulatory elements, marked by specific chromatin states and transcription factor occupancy, interact within the three-dimensional nuclear space to control precise spatiotemporal gene expression patterns^6^. While large-scale consortia like ENCODE^7^ and Roadmap Epigenomic^8^ have generated vast multimodal datasets—including gene expression, histone modifications, and chromosome conformation—a fundamental challenge persists: integrating these disparate data types into a unified, predictive, and generalizable model of genomic regulation. Such a unified model, if realized, is expected to provide important insight into the regulatory relation among the different regulatory mechanisms and thus provide a deep understanding of the determination of the cellular states and their response to stimuli.

Significant strides have been made in generative genomic modeling. Pioneering models such as Enformer^9^, Borzoi^10^ and AlphaGenome^11^ demonstrated that deep learning architectures trained on DNA sequence alone can predict a variety of cell-type-specific chromatin^12–15^ and expression profiles^16–20^, effectively learning a generalizable regulatory code from sequence context. However, by relying solely on sequence, these models are inherently constrained to predicting average profiles across the cell types in their training data; they cannot be dynamically conditioned on a specific cellular context provided by a user, limiting their utility for personalized prediction or for studying cell types absent from the training distribution. A separate class of models^21–31^, including Enformer-celltyping^24^, EPCOT^23,30^, GET^25^, and C.Origami^26^, addressed this task by incorporating cell-type-defining signals—such as (sc)ATAC-seq, DNase-seq or ChIP-seq profiles—as direct model inputs alongside DNA sequence. While this strategy enables cell-type-specific prediction, these models are often specialized for a single output modality (e.g., gene expression for GET, and chromatin contacts for C.Origami) and may suffer from pattern loss for samples drastically differ from the training cells. Concurrently, large language models (LLMs) adapted for genomics^32^, such as Evo^33^ and Nucleotide Transformer^34^, have shown remarkable promise in learning biological semantics from tokenized DNA sequences across scales. Yet, as purely sequence-based models, they share the limitation of being unable to ingest epigenetic context to condition their outputs on a specific cell state, separating their powerful pattern recognition from the mechanistic, context-aware simulation of the regulatory genome^35^.

Here, we introduce GenoME, a generative model designed to overcome these limitations through a novel Mixture of Experts (MoE) architecture^36^. GenoME integrates a 1-Mb DNA sequence with a corresponding cell-type-specific ATAC-seq or DNase-seq signal to generate a unified, multiscale profile encompassing epigenomics, transcriptomics, and 3D conformation at their native resolutions. Compared to previous approaches, the MoE framework allows the model to learn specialized, reusable “expert” functions for different regulatory concepts, facilitating robust generalization to new cellular contexts unseen during training. The built-in *in silico* perturbation framework also allows causal interrogation of regulatory functions by manipulating the input chromatin accessibility profile or DNA sequence. By unifying prediction, generalization, and perturbation within a single model, GenoME thus provides a versatile platform for exploring the causal logic of the multiscale regulatory genome, with potential applications in functional and personalized genetics.

## Results

### GenoME Enables High-Fidelity, Cell-Type-Specific Multimodal Characterization

GenoME is a Mixture of Experts (MoE)-based generative model designed for the cross-cell-type, multiscale prediction and perturbation of genomic profiles. For a given 1-Mb locus, the model integrates the underlying DNA sequence with a cell-type-specific ATAC-seq profile to generate a unified set of multimodal predictions. The tasks considered here include gene expression (total RNA-seq), transcription factor binding (TF ChIP-seq), a suite of histone modifications (Histone ChIP-seq), and 3D chromatin conformation (Hi-C contact maps), each at its native resolution (**Fig. 1a**).

**Fig. 1.**
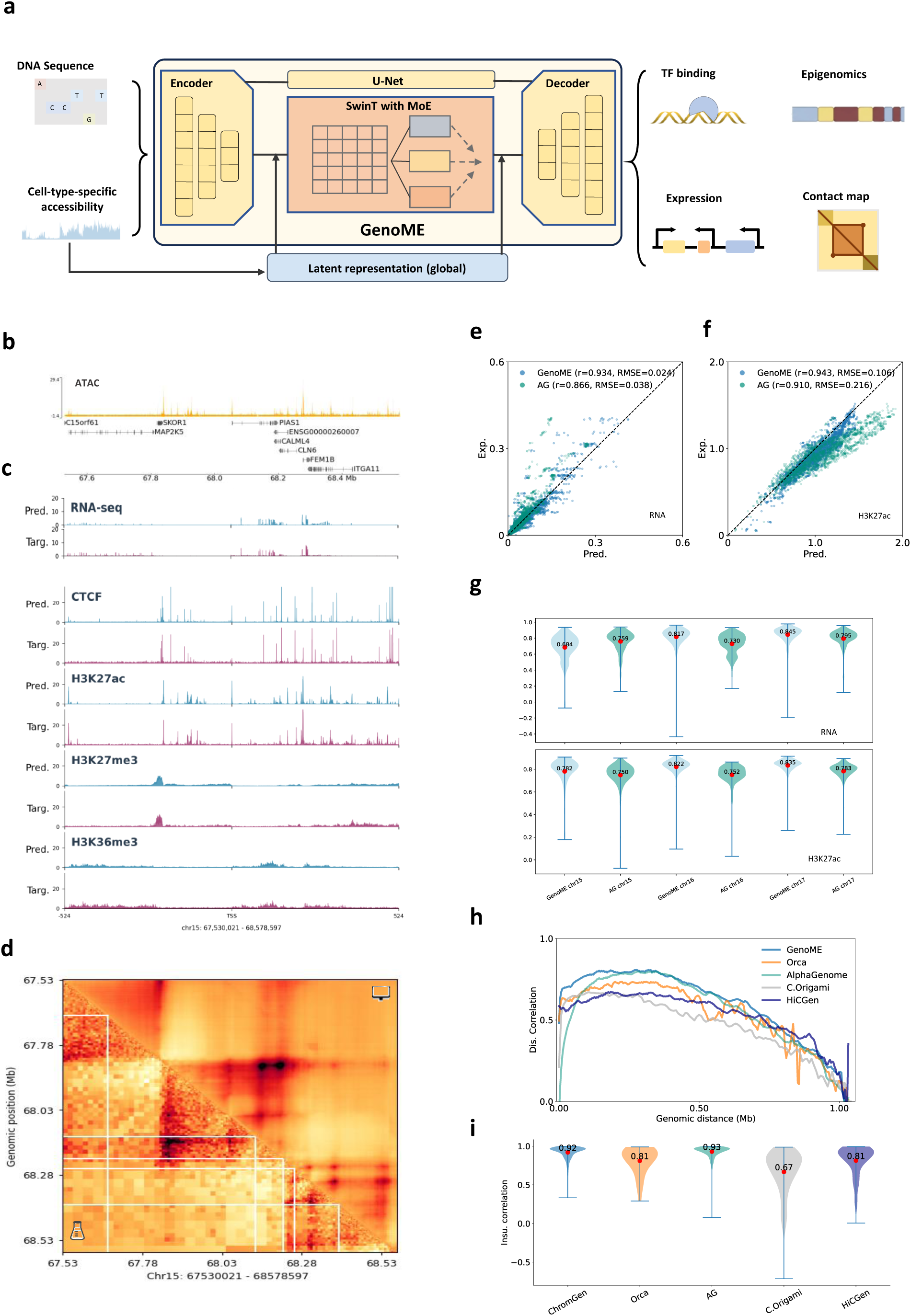
GenoME enables cell-type specific multi-modal characterization of the genome. a. With 1-Mb DNA sequence as well as cell-type specific ATAC-seq signals as inputs, GenoME enables predictions of epigenomics, gene expression, activities of cis-regulatory elements, TF bindings, and contact maps. b-d. exemplified 1-Mb (b) ATAC-seq inputs (blue), (c) the corresponding predictions (red) and targets (gray) of total RNA-seq at 1bp resolution, ChIP-seq (including CTCF TF-ChIP seq, H3K27ac, H3K27me3, and H3K36me3 Histone ChIP-seq) at 128bp resolution and (d) Hi-C maps at 2048bp resolution, for a selected region for heart right ventricle (male, 69 years). e-g. 2D comparisons of GenoME (blue) and AlphaGenome (green) model predictions with experimental targets for track-average (e) total RNA-seq signals at 1 bp and (f) H3K27ac Histone ChIP-seq at 128 bp for right lobe of liver, female, 47 years. (g) Comparisons of corresponding bin-wise correlations of model predictions with measurements. h,i. Comparisons of (h) stratified distance correlations and (i) insulation correlations of predicted Hi-C matrices determined by GenoME, Orca, AlphaGenome, C.Origami and HiCGen with experimental targets. All predicted and experimental maps were reshaped and/or truncated to a 1-Mb window at 4-kb resolution. Correlations were measured at test sets of the training cell type for each model (HCT116 for AlphaGenome and GenoME, HFF for Orca, IMR90 for C.Origami, and GM12878 for HiCGen).

As an example, we assessed the model’s ability to recapitulate experimental profiles in a held-out genomic region from human heart ventricle (**Fig. 1b-c**). Predictions closely matched experimental targets across all modalities, capturing fine-scale features such as sharp RNA-seq peaks at 1-bp resolution and complex histone modification patterns at 128-bp resolution (**Fig. 1c**). The model also successfully predicted higher-order chromatin architecture, generating Hi-C maps that reproduced characteristic domain structures and specific looping interactions **(Fig. 1d**).

Quantitative benchmarking confirmed the high fidelity of these predictions. GenoME achieved correlations with experimental data that were comparable to or surpassed those of baseline models. For example, in selected chromosomes from an adult liver donor, GenoME’s predicted H3K27ac and RNA–seq signals showed stronger alignment with experimental targets than did predictions from AlphaGenome (**Fig. 1e-g**). Note that AlphaGenome, a sequence–based model, is inherently constrained to generating average profiles for each tissue or cell type present in its training data. When predicting Hi–C maps on a held–out chromosome at 4–kb resolution, GenoME yielded superior distance–stratified and insulation correlations relative to specialized models including Orca and C.Origami (**Fig. 1h-i**). HiCGen exhibited an advantage in predicting long–range contacts (>0.7 Mb), likely because it incorporates genomic context beyond the 1–Mb input window.

Supplementary Figs. 2-7 present a more comprehensive benchmarking analysis, encompassing genome–wide correlation assessments across multiple modalities, quantitative evaluations across diverse chromosomes and cell types, and comparisons with recent models. In these analyses, GenoME-basic (the basic version of GenoME) consistently matched or exceeded the performance of established models for nearly all profiled genomic features—particularly features directly tied to gene regulation, such as H3K27ac and chromatin contact maps—underscoring its integrated design for capturing regulatory grammar.

### GenoME Generalizes to Unseen Cellular Contexts and Captures Inter-Individual Variation

Whereas foundational sequence–based models such as AlphaGenome learn to predict a single, averaged regulatory profile per tissue type from DNA sequence alone, GenoME is explicitly designed to condition its predictions on a provided, donor–specific epigenetic state. This capability is demonstrated through predictions of RNA–seq profiles for individual donors in tissues such as liver and psoas muscle (**Fig. 2a-b**). Compared with the average signals predicted by AlphaGenome for each tissue type, our model effectively tailors its predictions to the donor–specific ATAC–seq input (Supplementary Figs. 2-3).

**Fig. 2.**
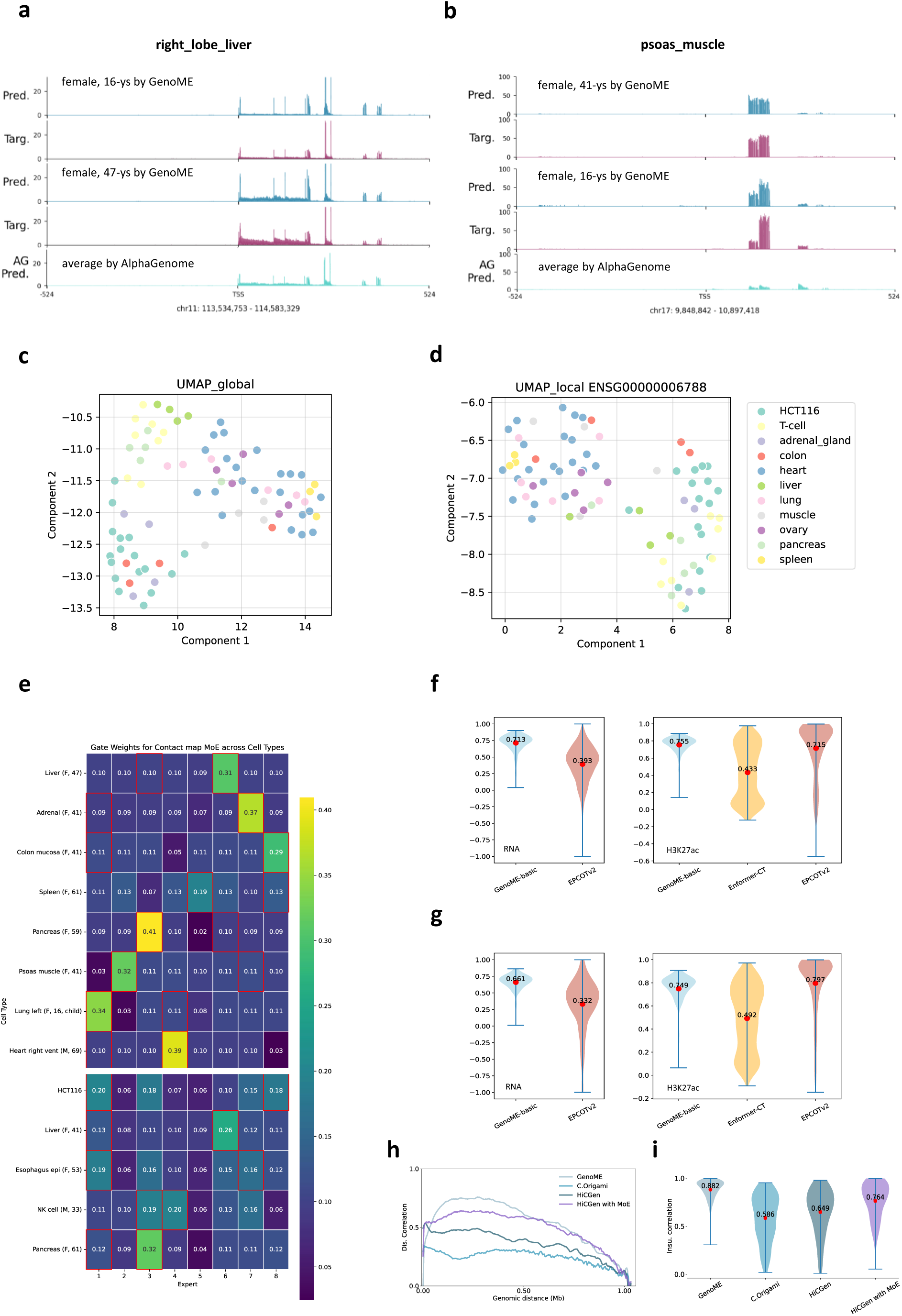
GenoME captures personalized genomic features via de novo cross-celltype predictions. a,b. Comparisons of selected genome regions with distinct expression profiles between different individuals for (a) right lobe liver and (b) psoas muscle tissue. The predicted RNA-seq profiles (blue) show good consistency with experimental results (red), while the tissue-average profile is predicted by AlphaGenome (green). c,d. UMAP projection of the (c) global embeddings of representative chromatin accessibility and (d) local embeddings of 1-Mb ATAC-seq signals centered around MYH13. Input ATAC-seq profiles are colored by their broad tissue type. Both global and local cell type embedding shows co-localization of similar cell types, but local UMAPs obtained from different input genome regions may vary greatly. e. Heat maps of mean gating weights respectively for conact map MoE layers across the test set chromosomes, for training (upper panel) as well as unseen (lower panel) cell types. Red box indicates the selected experts with largest output weights. f,g. Comparisons of bin-wise correlations of total RNA-seq and H3K27ac ChIP-seq tracks of *de novo* predictions of unseen samples, (f) right lobe liver female 41-ys and (g) esophagus squamous epithelium female 53-ys, for GenoME, Enformer-Celltyping and EPCOT2. h,i. metrics of de novo predictions of an unseen cell line, HCT116, in terms of (h) distance stratified correlations and (i) insulation correlations of Hi-C maps. All predictions were measured by GenoME-basic. Predictions of Hi-C maps were compared with C.Origami trained on IMR90, HiCGen trained on GM12878, and HiCGen-MoE trained on GM12878, IMR90, sigmoid colon, PANC-1 and prostate gland. With the help of MoE layers and multi-modal predicting heads, GenoME achieves the best performance on these tasks.

Analysis of the learned representations reveals the basis for this generalization ability. UMAP visualizations of both global and local (e.g., centered on the MYH13 locus, as in **Fig. 2b**) cell embeddings show that GenoME organizes cellular states in a biologically meaningful latent space, where clustering is driven primarily by tissue of origin (**Fig. 2c-d**). The context–dependent variation in local embeddings and the clear separation of certain donors from the same tissue type (e.g., liver and psoas muscle) in **Fig. 2d** indicate that the model captures subtle, region–specific differences in cellular state, thereby enhancing its adaptability to novel inputs. **Fig. 2e** presents heatmaps of the mean gating weights for the contact–map MoE layer across training and unseen cell types. Although certain experts are reused across cell types, both their activation levels and the combination of engaged experts are dynamically modulated by cellular context. Importantly, the gating network generalizes to unseen contexts: for example, the unseen pancreas sample (F, 61) exhibits an expert usage pattern highly similar to that of the training pancreas sample (F, 59). HCT116 shows an expert usage profile broadly comparable to that of colon mucosa (a training tissue) but with different weight distributions. This structured reuse, rather than random assignment, explains why HCT116 can partially overlap with other cell types in the UMAP while still maintaining a distinct regulatory grammar.

This generalization capability was then rigorously quantified. For de novo prediction of RNA–seq and ChIP–seq profiles in unseen liver and esophagus samples, GenoME–basic outperformed recent cell–type–specific models, namely Enformer-Celltyping^24^ and EPCOT2^30^ (**Fig. 2f-g**). More strikingly, when predicting the 3D conformation of the completely unseen HCT116 cell line, GenoME surpassed models specifically designed for Hi–C prediction—including C.Origami and HiCGen—achieving superior distance–stratified and insulation correlations (**Fig. 2h-i**).

The robust cross-cell-type performance stems from GenoME’s integrated MoE architecture, which allows knowledge sharing across modalities and cellular contexts through specialized expert layers. While decoder MoE gating network capture cell-type identities, SwinT MoE experts specialize in genomic-region-level regulatory contexts (Supplementary Fig. 9). Consequently, the model can construct a coherent, multimodal regulatory state for a novel cell type using only its ATAC-seq profile as a guide. Together, these results establish GenoME as a powerful tool for inferring regulatory landscapes from a single epigenetic input, even for cellular contexts absent from the training data.

### GenoME Enables *in silico* Perturbation for Mechanistic Investigation

Beyond static prediction, a paramount utility of a generative model is its capacity to serve as an *in silico* experimental system for probing causal regulatory relationships. We first validated this framework against experimental data from genetically engineered HCT116 cell lines featuring auxin-induced degradation of key regulatory proteins: CDK7, POLR2A, CTCF, and RAD21. For the HBG2 locus, GenoME accurately predicted the consequent changes in both RNA-seq and CTCF ChIP-seq signals (**Fig. 3a-b**). Critically, it generalized to the unseen perturbation of CDK7 degradation, successfully predicting the resultant transcriptional upregulation. Furthermore, the model’s predicted Hi-C maps for each perturbation recapitulated specific structural alterations observed experimentally, including the severe disruption of topological boundaries following depletion of CTCF or RAD21 (**Fig. 3c**) —an outcome poorly characterized by non-MoE models (Supplementary Fig.4).

**Fig. 3.**
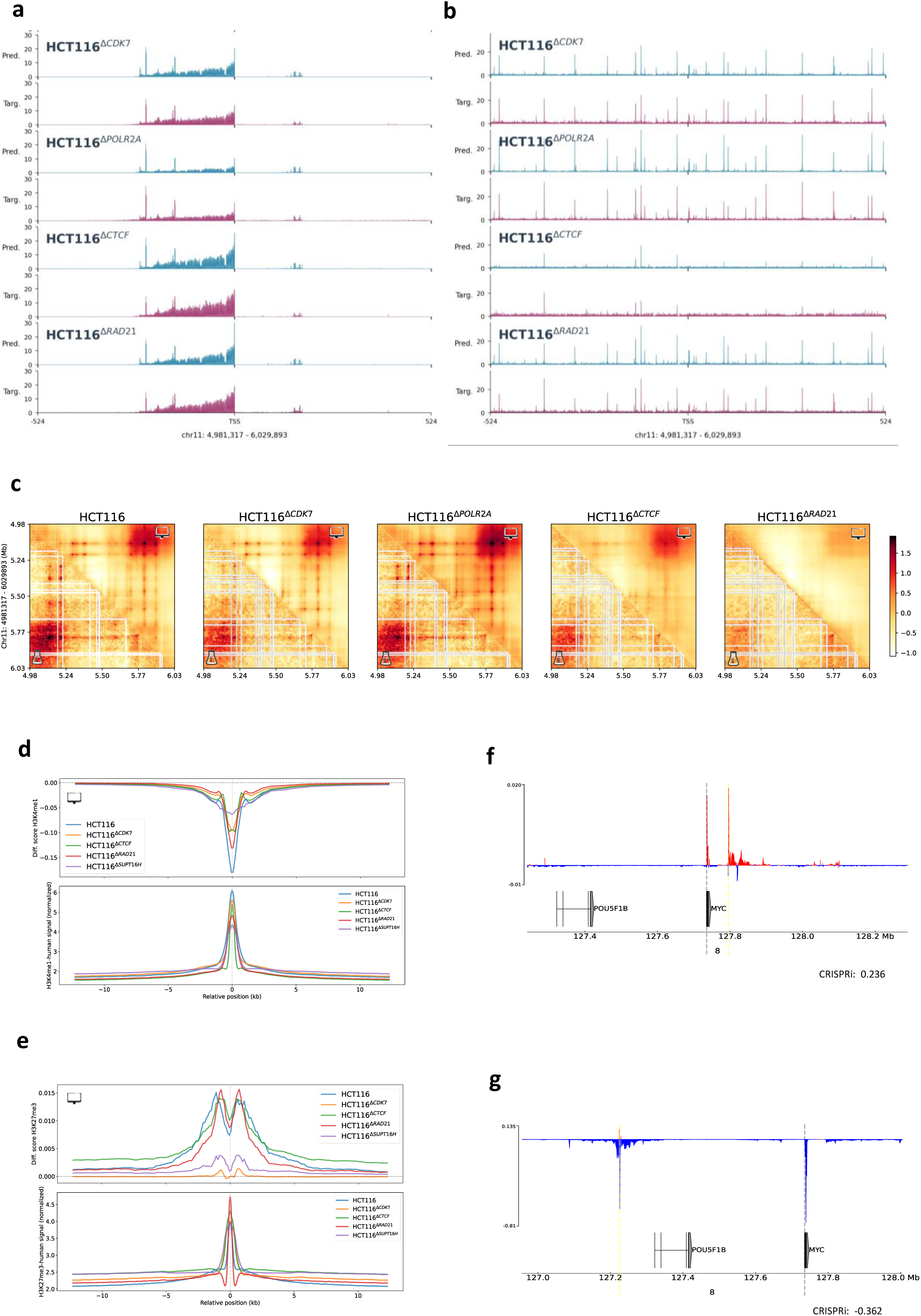
GenoME allows *in silico* perturbations through silencing/activating selected genome regions. a-c. Exemplified comparisons of (a) RNA-seq, (b) CTCF ChIP-seq and (c) contact maps predicted by GenoME-HCT116 with experimental signals of HBG2 region of HCT116 and its genetically modified counterparts HCT116^ΔCDK7^, HCT116^ΔPOLR2A^, HCT116^ΔCTCF^ and HCT116^ΔRAD21^ (which corresponding to Auxin-induced degradation of CDK7, POLR2A, CTCF and RAD21, respectively). HCT116^ΔCDK7^ was unseen during training. d,e. The average differential curves of (d) H3K4me1 and (e) H3K27me3 predicted by GenoME-HCT116 for 40,000 randomly selected genomic regions of HCT116 and its genetically modified counterparts, by silencing of 1-kb ATAC-seq peaks. Also presented are the average (d) H3K4me1 and (e) H3K27me3 ChIP-seq signals at overlapped regions with ATAC-seq peaks derived from ENCODE (lower panels). Wider impact range was observed for HCT116^ΔSUPT16H^ in H3K4me1 and HCT116^ΔCTCF^ in H3K27me3. f,g. The differential curves of total RNA-seq of HCT116 predicted by GenoME-HCT116 by *in silico* mutagenesis (ISM) of active cis-regulatory element of MYC nominated by Activity-by-Contact (ABC) and validated by CRISPRi-FlowFISH. Positive (f) and negative (g) changes at TSS and gene bodies indicate the silencer or enhancer role of the remote element respectively, both of which are consistent with effective changes in gene expression (lower right of each panel) determined by CRISPRi-FlowFISH.

**Fig. 4.**
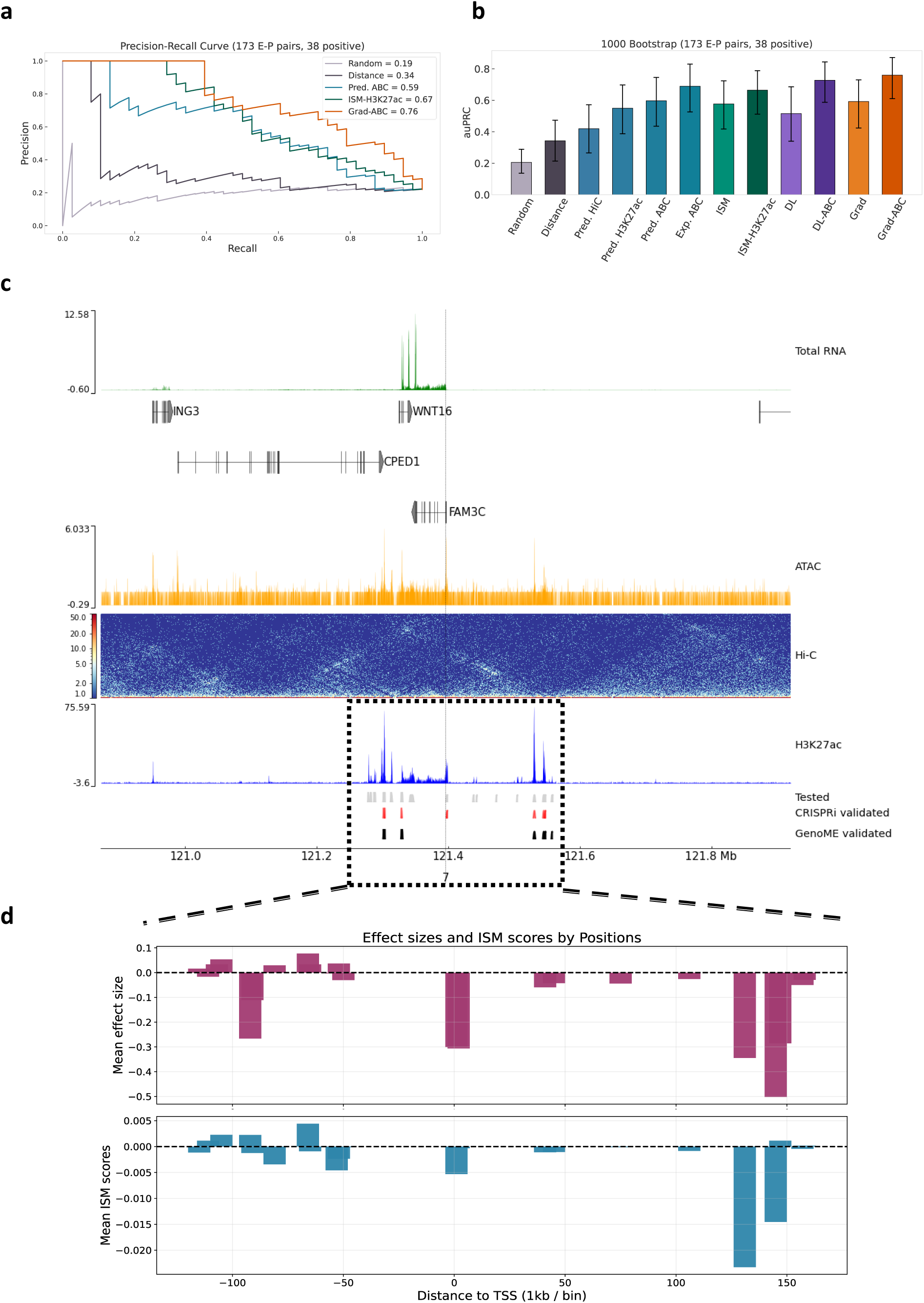
GenoME identifies activities of cis-regulatory elements. a. Precision-recall plot for classifiers of enhancer-promoter (E-P) pairs. Positive E-P pairs are those for which perturbation of the candidate enhancer significantly changes expression of the gene. The precision-recall curves represent the performance of the integrated gradient score of GenoME, *in silico* mutagenesis (ISM), DeepLIFT contributing score, predicted ABC score, and other assays on classifying 173 E-P pairs (with 38 positive ones) screened by CRISPRi-FlowFISH of HCT116. Predictions were measured by GenoME-HCT116. b. 1000 bootstap means of area under the precision-recall curve (auPRC) represent the performance of the E-P classifiers shown in (a) on classifying E-P pairs within 512-kb distance to the TSS. c. From top to bottom: the total RNA-seq, gene annotations, ATAC-seq, Hi-C contact, H3K27ac ChIP-seq, nominated enhancers for FAM3C tested and validated by CRISPRi-FlowFISH, and enhancers for FAM3C validated by Grad-ABC score, for 1-Mb locus centered at the TSS of FAM3C. The black dotted line marks the TSS of FAM3C . d. Bar plot illustrating the fractional change in gene expression resulting from (a )HCT116 CRISPRi-FlowFISH experiments and (b) ISM of each candidate regulator of FAM3C using GenoME within 200-kb distance to the TSS (black box shown in (c)). The sign of the scores indicate whether ISM the candidate enhancers within the bin would up-regulate (+) or down-regulate expression of the gene. A Pearson correlation coefficient of 0.53 was obtained from the scores of experimental and computational perturbations.

To systematically characterize the regulatory impact of local chromatin accessibility, we performed *in silico* silencing of 1–kb active ATAC–seq peaks across 40,000 randomly selected regions in wild–type and genetically perturbed HCT116 cells (Methods). Averaging the resulting differential predictions revealed distinct perturbation profiles. H3K4me1 modifications exhibited a slightly broader spread in SUPT16H–depleted cells (**Fig. 3d**), suggesting that altered nucleosome positioning reduces the precision of this activating mark. In contrast, H3K27me3 displayed a striking long–range effect, with differential signals propagating over hundreds of kilobases and accurately capturing Polycomb–mediated spreading (**Fig. 3e**). This repressive spread was further enhanced in cells lacking CTCF. In fact, CTCF actively opens chromatin and erases the H3K27me3 mark at its binding sites^37^. With the loss of CTCF, H3K27me3 can spread into the formerly insulated region, leading to a clear and widespread increase in the repressive mark^38^. By contrast, depletion of RAD21 produced no pronounced effect, in line with cohesin’s indirect role in H3K27me3 regulation via loop extrusion.

Experimental ChIP–seq data from regions overlapping the silenced ATAC–seq peaks validated these predictions: in ΔCTCF cells, H3K27me3 indeed invaded previously insulated regions, closely matching the average differential curve predicted by GenoME (**Fig. 3e**); likewise, H3K4me1 showed a broader perturbation spread in SUPT16H–depleted cells compared to other knockdowns (**Fig. 3d**), confirming the model’s perturbation framework.

Finally, we demonstrate the utility of this framework for functional annotation. Building upon the computational perturbations of selected enhancers act as context-specific silencers or activators, their functional roles can be directly validated through high-throughput screening experiments such as CRISPRa and CRISPRi^39–41^. For instance, *in silico* mutagenesis (ISM; see Methods) of two putative enhancers for the MYC gene, nominated by the Activity-by-Contact^41^ (ABC) model, respectively yielded specific positive or negative differential RNA-seq signals (**Fig. 3f-g**), which are consistent with the respective changes in effective gene expression (i.e. 0.236 and −0.362) determined in CRISPRi experiment. Collectively, GenoME establishes a powerful platform not only for forecasting experimental outcomes but also for large-scale, causal exploration of regulatory logic.

### GenoME Identifies Functional Enhancer-Promoter Connections

A central challenge in genomics is the functional annotation of non-coding sequences and the accurate linkage of distal cis-regulatory elements (CREs) to their target genes. To quantify the predicted regulatory influence of a candidate enhancer, we derived a novel “Gradient-ABC” score that multiplies the model’s integrated gradient by its predicted ABC score utilizing cell-type-specific ATAC-seq, predicted H3K27ac signal and Hi-C contact map (Methods).

Benchmarked against high-throughput CRISPRi-FlowFISH data in HCT116 cells—which provides a rigorous ground truth of 173 enhancer-promoter (E-P) pairs (38 validated positives)—the combinative score derived from the integrated gradient contribution, ATAC-seq inputs, the predicted H3K27ac and contact maps (i.e. Grad-ABC score) achieved superior precision-recall performance compared to the established ABC score, *in silico* mutagenesis (ISM) and DeepLIFT^42^ score (**Fig. 4a**). This advantage was robust, as confirmed by bootstrap analysis of the area under the precision-recall curve (auPRC) (**Fig. 4b**), and generalized effectively to the unseen K562 cell line (Supplementary Fig.5), demonstrating the transferability of the learned regulatory principles.

**Fig. 5.**
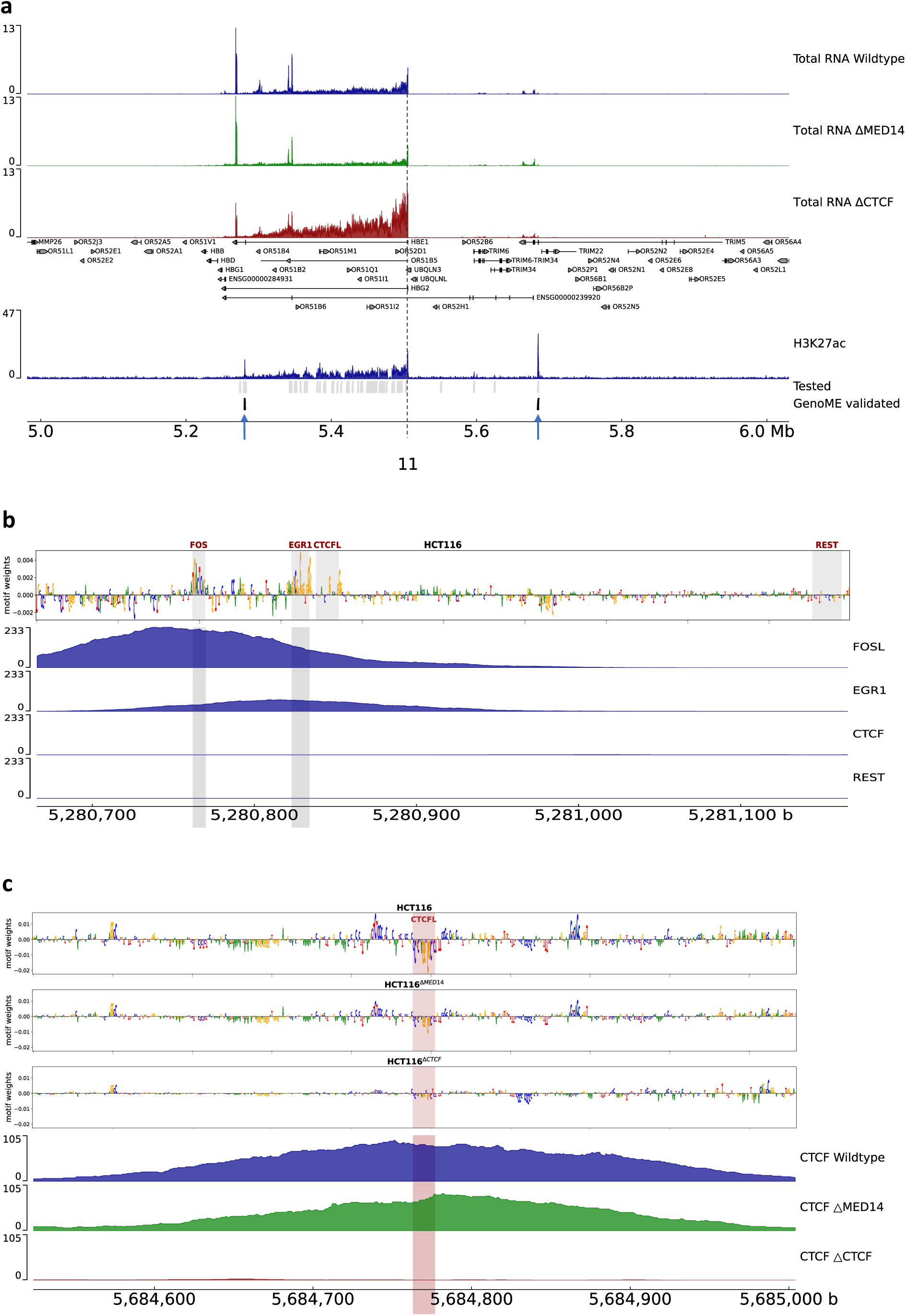
GenoME predicts transcription factor motifs at cell-type-specific enhancers. a. From top to bottom: the total RNA-seq, gene annotations, H3K27ac ChIP-seq, nominated enhancers for HBG2 tested and validated by Grad-ABC score, for 1-Mb locus centered at the HBG2 TSS of HCT116, HCT116^ΔMED14^ and HCT116^ΔCTCF^. The black dotted line marks the TSS of HBG2. b,c. The *in silico* mutagenesis (ISM) attribution score at (b) chr11:5,280,666-5,281,166 and (c) chr11:5,684,524-5,685,004 with top predicted enhancer activity (indicated by blue arrows shown in a), overlayed by the corresponding TF ChIP-seq data derived from ENCODE. All ISM inferences were measured by GenoME-HCT116. Sequences with high scores are highlighted in the boxes, with labels and arrows indicating the ID and names of the resembling known TF motifs in the JASPAR database.

To illustrate the quantitative predictive power of the model, we conducted a focused *in silico* perturbation analysis on regulators of the FAM3C gene in HCT116 (**Fig. 4c-d**). GenoME was used to predict the fractional change in FAM3C expression upon ISM of each candidate enhancer within a 512 kb window. The direction and magnitude of these ISM showed a clear positive correlation (Pearson *r* = 0.53) with the changes measured in the actuals CRISPRi-FlowFISH experiments (**Fig. 4d**). This concordance confirms that GenoME can not only classify functional connections but also quantitatively estimate the magnitude and direction (enhancing or silencing) of a regulatory element’s effect.

### GenoME Deciphers Cell-Type-Specific Transcription Factor Codes

The ability to infer the transcription factor (TF) code governing enhancer activity is a crucial step toward mechanistic understanding of transcriptional regulation. In **Fig. 5a** we applied GenoME to dissect the regulatory landscape of the HBG2 locus, where integrated gradient scores pinpointed distal enhancers active in HCT116 and its engineered variants (HCT116^ΔMED14^ and HCT116^ΔCTCF^).

*In silico* mutagenesis (ISM) of one active enhancer (chr11:5,280,666–5,281,166), performed with GenoME trained on HCT116 (GenoME-HCT116), generated nucleotide-resolution attribution maps (**Fig. 5b**). This analysis revealed specific, high-importance sequences matching binding motifs for known regulators, including FOS::JUN and EGR1. We reinforced this interpretation by overlaying ChIP-seq data: motif positions for FOSL and EGR1 coincided with ChIP-seq peaks of AP-1 family members (FOSL, JUND, SP1) and EGR1, respectively. In contrast, CTCFL and REST showed neither clear motif patterns nor strong ChIP-seq signals. These independent data confirm that GenoME correctly identifies activating motifs whose importance scales with expression output.

We further examined motif effects across HCT116 variants. As shown in **Fig. 5c**, CTCFL exerted a stronger repressive effect on predicted gene expression in HCT116 and HCT116^ΔMED14^ than in HCT116^ΔCTCF^. Notably, substantially stronger CTCF ChIP–seq signals were also observed in HCT116 and HCT116^ΔMED14^, indicating a context–dependent interplay between CTCFL–mediated repression and CTCF occupancy. Similar correlations between motif enrichment and TF peaks were obtained for additional loci (Supplementary Fig. 11).

## Discussion

A model that can accurately simulate the multiscale, cell-type-specific genome from first principles would dramatically accelerate our ability to interpret non-coding genetic variation, predict the consequences of genomic perturbations, clarify the relation between different epigenetic marks, and decode the logic of cellular identity. Here we present GenoME, a multimodal generative model that successfully unifies the prediction of genomic features from DNA sequence and chromatin accessibility across scales and cell types. By leveraging a Mixture of Experts architecture, GenoME moves beyond single-task or single-cell-type models to offer a versatile, all-in-one platform for genomic analysis. The MoE architecture learns reusable regulatory “building blocks” rather than strictly disjoint cell-type representations. The encoder (SwinT) MoE layers specialize in genomic-region-level contexts (e.g., forest/prairie compartments^43^), while the decoder MoE layers exhibit cell-type-specific routing patterns. Our results demonstrate its capabilities in multiscale and multi-modal characterization and cross-cell-type generalization.

While integration is a strength, we note that adding output modalities does not uniformly improve performance across all tasks. For instance, incorporating a splice junction prediction head slightly reduced accuracy for other tracks (Supplementary Fig. 7). This observation aligns with ablation studies in other generative models, such as AlphaGenome, where adding certain task heads had negligible effects on unrelated outputs. This phenomenon likely reflects the inherent tension between shared and task-specific representations within a unified model.

Adding a complex prediction target, such as splice junctions, may introduce a supervisory signal whose optimal feature space partially conflicts with or dilutes features critical for other tasks.

Beyond static prediction, GenoME serves as a versatile *in silico* experimental platform and provides a direct path from functional prediction to mechanistic insight. The integrated gradient scores derived from the model outperformed the established ABC model in classifying functional enhancer-gene pairs from CRISPRi screens. When combined with *in silico* mutagenesis, this approach successfully identifies transcription factor motifs driving activity at predicted enhancers, connecting sequence syntax to regulatory function in a cell-type-specific manner.

Despite its strengths, GenoME has limitations that point toward future development. Its current context window of 1 Mb, while sufficient for many locus-specific investigations, prevents analysis of long-range interactions or whole-chromosome organization. Furthermore, the model currently treats chromatin accessibility as a static, condition-defining input. Future iterations could incorporate time-series or stimulus-responsive data to model dynamic regulatory processes. Integrating genetic variation as an explicit input would allow direct prediction of the multiscale functional consequences of SNPs and other variants, bridging closer to personalized genomics.

Finally, to improve efficiency and accuracy with even longer sequence lengths, more sophisticated architectural designs—such as hybrid models incorporating Mamba ^44^ or Hyena ^45,46^—are promising directions.

In conclusion, GenoME represents a unified, cell-type-specific model of the multiscale regulatory genome. Its ability to generalize across contexts and simulate perturbations bridges the gap between correlative observation and causal understanding. We envision it becoming a platform for interpreting non-coding genomes, prioritizing functional elements, and ultimately enhancing our ability to decipher the complex regulatory code of life^47^.

## Materials and Methods

### Data preprocessing

**Reference Genome:** the genomic sequences for training the model were derived from human GRCh38/hg38 reference genome. Gene and transcript annotations used for evaluations were based on the GTF V47 annotations derived from the GENCODE data portal (https://www.gencodegenes.org/). We performed five-channel one-hot encoding of the genomic bases ‘ACGT’ and the unknown base ‘N’. For sequence intervals that extended beyond chromosomal boundaries, we used ’N’ characters for padding to maintain the consistency of input length.

**ATAC-seq Data:** the sequencing data for ATAC-seq experiments were obtained from the ENCODE data portal (https://www.encodeproject.org/). For quality control of the ATAC-seq data as model input, we stringently selected tracks that met the following conditions:

1. The data contained neither ‘ERROR’ level ENCODE audits, nor ‘NOT_COMPLIANT’ level audits related to insufficient read depth, insufficient number of reproducible peaks, or severe bottlenecking;
2. The experiments were performed either on Illumina NovaSeq 6000 or Illumina HiSeq 4000 platform;
3. The tracks had a library fragment size of 200-7000 nt and a read length of 100 nt.

For each ATAC-seq experiment, we first downloaded the raw alignment (BAM) files for individual replicates. The BAM files were merged and subsampled down to 50M reads (samples with reads less than 50M were removed). Next, the down-sampled BAM files were converted into bigWig files following the procedure of ChromBPNet^48^. The final outputs shall preserve the base-resolution nature of the data and facilitate decoupling enzyme cut bias (by Tn5) from true signal by applying appropriate read shifts. By taking the above quality check, we finally obtained 63 groups of ATAC-seq tracks in tissues, 9 groups in primary cells and 17 groups in cell lines. Details of all accession number are listed in Supplementary Table 1-3. To improve numerical stability, we transformed each ATAC-seq track signal as follows:

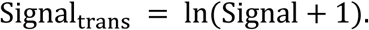

The global chromatin accessibility corresponds to the chromatin accessibility for 3 kb around the transcriptional start site of 1216 marker genes nominated by PangloaDB^49^ was also labeled as cell-type features feed to the Mixture of Experts (MoE) layer for better gating during training. These global signals were averaged at 250-bp resolution and then compressed into an input in length of 14,592, providing a cell-type-defining context vector. See Supplementary Table 4 for the full list of these marker genes.

**RNA-seq Data:** the sequencing data for total RNA-seq experiments were also obtained from ENCODE. The selected tracks met the following conditions:

1. the data contained neither ‘ERROR’ level ENCODE audits, nor ‘NOT_COMPLIANT’ level audits related to insufficient read length;
2. the bio-sample summary and term of the track corresponded to one of the selected ATAC-seq tracks;
3. the tracks had a minimum library fragment size of 200 nt and a minimum read length of 100 nt.
4. the data were released not earlier than 2019.

We selected the stared bigWig files with name ‘minus strand signal of unique reads’ and ‘plus strand signal of unique reads’. If multiple bigWig files per experiment and output type correspond to a single ATAC-seq track, the one with less severe audit flags and latest processing date was chosen. Details of all accession number are listed in Supplementary Table 1-3. Each individual RNA-seq signal track was normalized to represent a total of 100 million reads and then divided by the mean of their non-zero values. To dampen extremely high signals, we further applied scaling and clipping transformation to each track as follows:

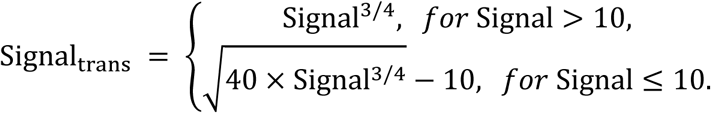

**ChIP-seq Data:** for TF ChIP-seq (CTCF), Histone ChIP-seq and Mint-ChIP-seq signals (H3K27ac, H3K27me3, H3K4me1, H3K4me3, H3K9me3, H3K36me3), we selected fold change over control bigwig files from ENCODE. through the ChIP-Route. The selected tracks met the following conditions:

1. the data contained neither ‘ERROR’ level ENCODE audits, nor ‘NOT_COMPLIANT’ level audits related to severe bottlenecking;
2. the bio-sample summary and term of the track corresponded to one of the selected ATAC-seq tracks;
3. the tracks had a minimum mapped read length of 50 nt.
4. the data were release not before than 2019.

Similarly, If multiple bigWig files per experiment type correspond to a single ATAC-seq track, the one with less severe audit flags and latest processing date was chosen. Details of all accessing numbers are listed in Supplementary Table 1-3. The tracks were also respectively divided by the mean of their non-zero values and clipped at 10, similarly to the handling for RNA-seq signals (but without the 3/4 scaling).

**Hi-C Data:** The intact Hi-C data were derived from the call sets from the ENCODE portal with identifiers listed in Supplementary Table 1-3. All data were mapped to hg38 in cool format ^50^. Raichu normalization^51^ with default parameters were performed to the unbalanced data to remove biases. Bins with fewer than 10 nonzero counts were removed from the balanced matrix. Adaptive coarse graining procedures ^52^ were applied to the contact matrices to smooth the low-coverage areas at 1-kb-resolution. The average contact frequency as a function of genomic distance (i.e. genomic-distance-based background) was determined by cooltools^53^. Each

Hi-C data was then normalized as the log-fold contact map over the background as follows:

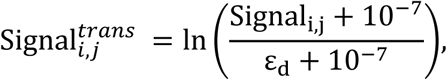

with ε_d_ representing the averaged background at genomic distance *d*=abs(i-j).

**Splice Junctions:** The splice junctions were derived from the BAM files of the corresponding total RNA-seq datasets. By referring to GTF V47 annotations derived from the GENCODE, each BAM file was processed by samtools and then realigned with STAR^54^ (Splice-aware Transcriptome Aligner) to discover splice junctions. The alignment was performed with a minimum splice junction overhang of 8 base pairs, allowing a maximum of 20 alignments for multi-mapping reads. The minimum and maximum allowed intron length were set to 20 and 10^6^, respectively.

The output junctions were only retained if its strand information is specific and its read count is greater than 3. The retained junction counts for each sample were normalized to 1 million total filtered junction reads for each sample. For numerical stability, the retained junction counts were first clipped at 1000 and then divided by their mean values for each sample.

The splice sites were derived from the AlphaGenome^11^ output head of the splice site classification. For any giving input genome interval, the splice sites (i.e. doners and acceptors) were determined respectively as the top 512 sites of highest probability from the output of the splice site classification for each strand. The splice junctions were then represented as a 512×512 matrix corresponding to the donor-acceptor pairs for each strand. Each element of the matrix was filled by the corresponding experimental counts for each donor-acceptor pair (padded with 0 otherwise).

### Model architecture

GenoME takes 1-Mb sequence and genomic features (ATAC-seq or DNase-seq) as inputs, and down-samples the represented length from single nucleotides down to 128 bp for each bin by deploying seven blocks of 1D convolutional neural network (1D-CNN). The 1D-CNN module is implemented mainly for reducing the spatial dimension and adding channels of each token.

The embeddings of the sequence and genomic features at 128 bp are processed based on the Swin-transformer^55,56^ (SwinT) and Mamba ^44^ architecture. The transformer tower consists of five consecutive SwinT blocks with a window size _s*w*in_=256. The token length is halved after each SwinT block, and the channel dimension is doubled after the third SwinT block (Supplementary Fig. 1). The transformer tower outputs embeddings at 128, 256, 512, 1024 and 2048-bp resolutions, processed with 4, 2, 2, 2, and 2 Swin-attention layers, respectively.

For the initial two Swin-attention layers, we have a mixture of experts (MoE) layer integrated within each Transformer block’s feedforward network (Supplementary Fig. 1). The MoE layer consists of one shared expert and five specialist experts. Each expert is implemented as a two-layer neural network with GELU activations and dropout. The gating network determines expert selection by processing both local sequence embeddings and a global cell-type context vector. This global context vector, linearly projected to 128 dimensions, enables the model to condition expert selection on specific cellular states. For each input token, the gating network computes logits for the eight cell-type experts, applies SoftMax with temperature τ annealing from 0.8 to 0.5 with decay rate 0.95, and selects the top-2 experts. Two auxiliary losses ensure balanced expert utilization: (1) a z-loss (coefficient 0.0001 with decay rate 0.95) penalizing large logit magnitudes for numerical stability, and (2) a load-balancing loss (coefficient 0.001 with decay rate 0.95) encouraging equitable expert usage by penalizing high variance in gating weights. This design enables specialized expert functions to develop for different regulatory patterns while maintaining the flexibility to recombine them for novel cellular contexts using only the ATAC-seq-derived context vector.

The decoder for contact map is of a 2D convolutional neural network (2D-CNN). The module takes embeddings at 2048-bp resolution generated by SwinT as input, generating contact map with a window size of 512×512 matrix. 2D-CNN starts with a MoE gating network with one shared and eight specialist experts, followed by five dilated residual blocks. The final predictions of the contact map are generated by a head block with a GELU activation and linear transformation.

The decoder for 1-D signals employs up-sampling and a U-Net-style skip connections to the encoder at 1024, 512, 256, 128, 64, 32, 16, 8, 4, and 2-bp resolutions (Supplementary Fig. 1). The output heads for ChIP-seq and RNA-seq tracks respectively takes the 128-bp and 1-bp embeddings (after U-Net up-sampling) as inputs. Each output head contains a MoE gating network with one shared and eight specialist experts, a convolutional layer, a linear transformation and Softplus activation.

The output head for splice junctions predicts counts for potential junctions between donor and acceptor sites, taking the 1-bp resolution embeddings as inputs. First, a linear layer projects the embeddings to an intermediate dimension of 512 channels. The positions for 512 donor and acceptor sites (for each strand) of highest probability from AlphaGenome’s prediction are embedded through the RoPE formulation. Finally, the predicted counts for each junction are obtained via an inner product between the processed donor and acceptor embeddings, followed by a Softplus activation.

### Training

The model was trained on a GPU cluster with 8 NVIDIA Tesla A100 80GB GPUs. The data-loader generates training data, including sequence, ATAC-seq and total RNA-seq signals at 1-bp resolution, ChIP-seq signals at 128-bp resolution and Hi-C maps at 2-kb resolution. DNA sequence and ATAC-seq signals in length of 1 Mb were implemented as model inputs. The data were obtained by uniformly sampling the annotated gene regions from training chromosomes. Chromosomes were split into training, validation and test set based on the Chromosome ID. The entire chr10 and chr15 were respectively assigned to the validation and test set, while the rest were assigned to the training set. To enhance prediction accuracy, non-readable regions (including telomeres or centromeres) were masked. To reduce overfitting during training, each sampled interval was shifted by a randomly sampled distance from −8192 to +8192 bp. We also replaced about half of the forward-sequences with their reverse-complement counterparts during the training process. Gaussian noise with zero mean and 0.1 standard deviation was added to input signals.

For comparison, we trained three models, i.e. GenoME-basic, GenoME-finetuned (GenoME-ft) and GenoME-HCT116, each with specific training cell types and targets. GenoME-basic was trained on 8 selected tissues (Supplementary Table 5) for 100 epochs (without early-stopping). The training data from the selected tissue donors contained full targets (i.e. 2 RNA-seq tracks, 7 ChIP-seq tracks and 1 Hi-C map for each sample).

GenoME-ft was first pretrained on 64 groups of ATAC-seq tracks in tissues and 9 groups in primary cells for 100 epochs (without early-stopping), and then finetuned with 8 selected tissues same to GenoME-basic for another 100 epochs. GenoME-HCT116 was trained on 15 groups of original and genetically perturbed HCT116 samples (Supplementary Table 5) for 100 epochs (without early-stopping). The batch size was set to B=16 for pretraining, and B=8 for GenoME-basic, GenoME-HCT116 and finetuning GenoME-ft. The pretraining was performed on 8 A100 GPUs (2 batches per GPU). The other tasks used 4 GPUs, requiring approximately 80 hours each. Unless otherwise specified, the predictions and metrics in the main text were generated by the GenoME-basic model.

The training loss for ChIP-seq and RNA-seq tracks is calculated as a weighted sum of Poisson and Multinomial negative log-likelihoods, which can be written as:

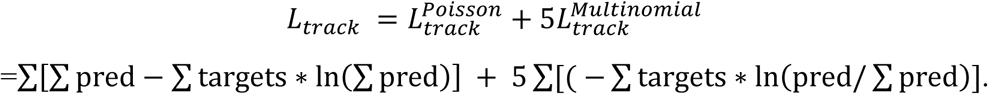

For numerical stability, the relevant sequence axis is divided into 8 equal segments before loss calculation for each track. Moreover, coverage at highly expressed exons may be several orders of magnitude higher than those of background for RNA-seq tracks. Therefore, at positions where the RNA-seq signal exceeds ten times the average, we double the weight in the loss calculation. To promotes tissue-specific expression patterns^57^, we also add an auxiliary gene-level loss contributes to the model’s total training loss with an overall weight of 0.1 for RNA-seq targets, which is calculated as the Poisson negative log-likelihoods of the total counts for each gene.

For predicting splice junctions, the loss is a weighted combination of the cross-entropy and Poisson components, which reads as:

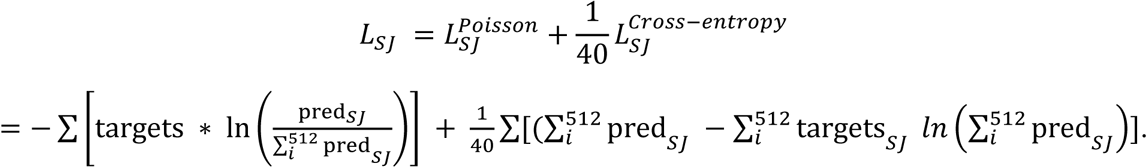

The loss is performed independently for positive and negative strand predictions using their respective splice sites.

Contact matrices at lower resolutions were obtained by average pooling the original Hi-C data. Contact matrices were down-sampled to meet the prediction windows at 2-kb resolution. The model was trained to predict the log-fold contact maps over genomic-distance-based background. The output matrices were compared with the experimental Hi-C map via a mean squared error (MSE) loss.

We trained the model with Adam optimizer, with a maximum learning rate of 2×10^-4^ and a weight decay of 10^-4^. The learning rate followed a schedule with a linear warmup from 0 to 2×10^-4^ over the first 20 epochs, followed by a Cosine annealing to 0 over the remaining 80 epochs.

### Evaluation

For cell-type-specific predictions, the model was provided with two inputs: (1) the 1-Mb reference DNA sequence from the target locus, and (2) the ATAC-seq profile from a cell type within or completely absent from the training data. No further fine-tuning on the target cell type was performed. The model’s predictions across all modalities were then compared to the experimental data from the specific cell type.

The performance of GenoME and comparator models was assessed using standard correlation metrics. For genomic tracks (RNA-seq, TF, and histone ChIP-seq), we computed the track-averaged and bin-wise Pearson correlation coefficient between the predicted and experimental signal at the native resolution of each track for the held-out test set (chr15). For unseen cell-types, we also computed whole-genome benchmarking for other chromosomes. For RNA-seq prediction, we also measured gene-level correlations by averaging counts within specific exon regions. For Hi-C contact maps, we calculated distance-stratified correlation by computing the correlation coefficient separately for pairs of genomic bins within 1 Mb intervals. We also evaluated insulation score correlation and observed-over-expected (O/E) correlation for the contact maps as described previously^27^. To evaluate predictive fidelity, we benchmarked GenoME against state-of-the-art models:

**AlphaGenome.** AlphaGenome is a unified sequence-based model offering multimodal predictions of gene expression, splicing patterns, chromatin features, and contact maps. As a purely sequence-based model, AlphaGenome outputs a single, tissue-averaged prediction per track and cannot be conditioned on a donor-specific epigenetic state. Prediction results for the identical 1-Mb DNA sequences from our test set were obtained via the official AlphaGenome API (https://www.alphagenomedocs.com/api). For benchmarking ChIP-seq and RNA-seq tracks in the right lobe of liver, we used the model output corresponding to the standard tissue identifier UBERON: 0001114. For benchmarking ChIP-seq and RNA-seq tracks in pancreas, we used the model output corresponding to the standard tissue identifier UBERON: 0001264. For benchmarking Contact map in HCT116, we used the model output corresponding to the EFO ID: 0002824.

**Enformer-Celltyping.** The ATAC-seq and six histone ChIP-seq experiments, along with their corresponding control BAM files, were downloaded from ENCODE and converted to the hg19 reference genome using liftOver to ensure compatibility with Enformer-Celltyping. The converted BAM files were then processed through the ENCODE standard ATAC-seq or ChIP-seq pipelines, with the number of fragments capped at 50 M to generate the p-value signal tracks. Using the resulting ATAC-seq signal track, we applied the model weights provided in Enformer-Celltyping to generate genome-wide predicted histone-mark profiles. We compiled the union of genomic intervals used during Enformer-Celltyping training across all cell types. Within each interval, we partitioned the sequence into 128-bp bins and computed the correlation coefficient between predicted and observed signal values. The results were compared against imputed EpiMap signal tracks for these samples. For the training-set evaluation, we selected two representative groups of samples (i.e.

ENCSR654UYP and ENCSR308HPZ). To evaluate the performance of Enformer-Celltyping on data not included in the training set, we selected one ENCODE sample (ENCSR516CPW) as an independent test case.

**EPCOT(v2).** The ChIP-seq data were preprocessed based on the EPCOT2 protocol. The ChIP-seq signals were downloaded from ENCODE in signal p-value format, followed by z-score normalization and trimming to the range (−2, 36), consistent with the EPCOT2 protocol. RNA-seq data were obtained from ENCODE with strand-specific separation. The ATAC-seq tracks were generated from BAM files and converted to RPGC format using bamCoverage. The default pretrained weights provided by EPCOT2 were used in benchmarking. Signals were predicted within a 500 kb window at the default 1 kb resolution. We took the exp(x) − 1 transformation for the RNA-seq tracks. For Hi-C data, we evaluated the O/E correlation with SCALE-normalization of the contact maps. Next, the target and predicted matrices were aggregated to 2-kb resolution. Based on the data preprocessing pipeline of EPCOT2, a Gaussian filter with a width of 2 was applied to smooth the experimental contact matrices. Subsequently, the experimental Hi-C data were trimmed to the range of −2 to 10, consistent with the preprocessing procedure of EPCOT2.

**GET.** Since GenoME metrics gene-level expression on held-out test chromosome, we derived leave-one chromosome-out benchmarking results of the general expression transformer (GET) from their paper^25^. As reported, an average Pearson correlation of 0.78 (minimum: 0.73, maximum: 0.84) was obtained on fetal astrocytes.

**Hi-C predictors.** We presented comparison with previously released Hi-C predictors AlphaGenome^11^, Orca^14^, C.Origami^26^ and HiCGen^27^. Since AlphaGenome was trained to predict a 1-Mb window of contacts, we restricted regions for comparison to 1-Mb blocks. If possible, we selected 1-Mb held-out genomic regions from cell types present in their respective training data or completely unseen cell types. For AlphaGenome, we chose chr15 predicted by the distilled model (which inevitably introduced data within training set) ; for Orca, we selected chr10 predicted by the model trained on HFF; for C.Origami, we used chr15 predicted by with the model trained on IMR90; for HiCGen, chr15 predicted by the model trained on GM12878 was used. We performed additional genomic-distance-based normalizations to the predicted results (and also the target maps) of C.Origami , which was already preprocessed to the training targets of AlphaGenome, Orca and HiCGen. To align with the 8-kb resolution of the prediction of C.Origami, we selected the 2-Mb window in Orca and HiCGen and cropped out the centering 1-Mb window for comparison. Since both AlphaGenome and Orca do not allow unseen cell-type predictions, benchmarking performance on *de novo* prediction of unseen cell types was compared only with HiCGen and C.Origami.

### *In silico* perturbation framework

A core capability of GenoME is its use as a predictive, *in silico* experimental system. The framework allows causal interrogation of regulatory logic by manipulating model inputs and observing the consequent multimodal predictions. Perturbations are applied to the inputs of a trained model, and the forward pass is recomputed to generate a new set of multiscale profiles. The differential between the perturbed and reference predictions quantifies the estimated effect. Three primary perturbation modes are implemented:

**Epigenetic signal activation/silencing.** The cell-type-defining ATAC-seq input profile is directly modified. This simulates experimental interventions that alter chromatin accessibility, such as CRISPR-mediated enhancer activation/silencing or the effects of protein degradation on chromatin state. Signal silencing is implemented by setting ATAC-seq signal values larger than ln(2) within a defined genomic window to ln(2). Signal activation is implemented by replacing signals with a boosted or exogenous profile (e.g., from another cell type).

**DNA sequence perturbation (*In silico* mutagenesis, ISM).** The one-hot encoded DNA sequence input is altered. This simulates the impact of genetic variants or targeted edits. For a given genomic window, every possible single-nucleotide substitution (A, C, G, T) is applied at each position sequentially. The change in a specific model output (e.g., gene expression log-counts) relative to the wild-type sequence is calculated as the attribution score for that position and nucleotide.

**DeepLIFT/SHAP.** We utilized via Captum^58^ for calculating nucleotide-specific DeepLIFT/SHAP scores. DeepLIFT contributions were averaged over 50 dinucleotide–shuffled baselines with ATAC signal set to its mean value. This process entailed generating 50 dinucleotide–shuffled variants of each sequence to serve as reference points. Subsequently, the importance scores obtained from DeepLIFT/SHAP^42^ for each sequence were combined with their respective one-hot-encoded matrices, yielding the final nucleotide contribution scores.

**Integrated gradient attribution.** To score the importance of specific input features (sequence or chromatin accessibility) for a given prediction, integrated gradients are computed. This provides a continuous attribution map across the input window, highlighting bases or accessible regions most critical for the model’s output. To simulate a continuous transition between cellular states, the ATAC-seq profile is morphed from a source state (e.g., HCT116 wild-type) to a target state (e.g., HCT116^ΔCTCF^) via linear interpolation across N=50 steps. For each intermediate step i, the input signal is computed as:

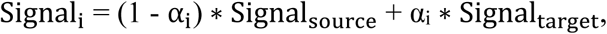

where α_i_ = i / (N-1). The model’s outputs (e.g., gene expression, H3K27me3, contact maps) are computed at each step, generating a trajectory of predicted consequences.

### Perturbation analysis

The general framework described above was applied to conduct specific causal analyses:

**Simulation of genetic perturbations**. The effects of protein degron experiments (e.g., auxin-induced degradation of CTCF, RAD21, CDK7, POLR2A) were modeled by providing GenoME with the ATAC-seq profile from the corresponding genetically modified cell line (HCT116^ΔCTCF^, etc.) as input, while keeping the reference DNA sequence unchanged.

**Regional epigenetic silencing/activation**. We first categorized 1-kb genomic fragments with at least 1 bp ATAC-seq signal values larger than ln(2) to active regions. Their ATAC-seq peaks were silenced or replaced with signals from an active state (e.g., a cancer enhancer profile applied to a normal tissue background). The mean absolute difference across all predicted modalities before and after perturbation defined an impact score. For instance, by silencing perturbations across 40,000 active regions, the average differential curve for histone marks (e.g., H3K27me3, H3K4me1) was calculated to quantify the genomic range and magnitude of epigenetic change. The differential curve was compared to the average ChIP-seq signals at overlapped regions with ATAC-seq peaks, which was calculated as the genome-average ChIP-seq signals with 30-kb window size centered at the same peak positions derived from ENCODE.

**Gradual epigenetic transition**. The cell-type modulation was used to trace the non-linear progression of outputs like H3K27me3 spreading or contact domain dissolution as the chromatin accessibility landscape was progressively transformed from a wild-type to a CTCF-depleted state.

### Enhancer-promoter pair prediction

We obtained enhancer–promoter (E-P) pairs from CRISPR-interference followed by FlowFISH^40,41,59^ (CRISPRi-FlowFISH), which perturbs enhancers and measures gene expression changes in HCT116 and K562 cells. For HCT116 experiment, we selected E-P pairs within 512 kb of the transcription start site (TSS), identifying 173 enhancer-promoter pairs with 38 active ones (Supplementary Table 6). The active enhancers significantly reduced gene expression (adjusted p-value < 0.05) with more than 80% power to detect a 25% effect size after perturbation. E-P pairs for K562 were derived from the CRISPRi dataset^54^.

We introduced different scores to prioritize E-P pairs. Candidate enhancers were derived from ENCODE annotations, which were defined by the activity-by-contact (ABC) method^41^. The ABC score for each E-P pair was calculated as the product of ATAC, H3K27ac signal and Hi-C contacts, while the predicted ABC score was calculated as the product of ATAC, predicted H3K27ac signal and Hi-C contacts. The *in silico* saturation mutagenesis (ISM) score was determined by setting each base pair in the enhancer region to the unknown nucleotide N in the vocabulary set and comparing against the wild-type magnitude of the corresponding gene expression. The DeepLIFT score was determined based on the 10-fold baseline with dinucleotide–shuffled sequences and ATAC signal setting to the mean value.The gradient scores were calculated as the integrated gradient (IG) from two types of baseline inputs within the selected enhancer regio: 1) uniform distribution of encoding DNA (i.e. 0.25 for each nucleotide) with true ATAC-seq signals, Grad_seq_, and 2) ATAC-seq signals setting to its mean value with true genomic sequence, Grad_atac_. The integral step was set to 50. A min-max normalization was next applied to both Grad_seq_ and Grad_atac_.The Grad-ABC score was calculated as the product of the model’s IG and the predicted ABC score. Note that the IG score we compute is highly targeted: for each enhancer--gene pair, the predicted RNA–seq track is reduced to a single scalar--the mean predicted RNA level over the selected gene interval (on specific strand). IG is then calculated with respect to this scalar and measured only within the enhancer coordinates of interest. Consequently, the gradient score reflects how changes in the sequence of enhancer and local chromatin features affect that specific gene expression. We also introduced other assays to score E-P pairs, including Hi-C contacts, H3K27ac signal, and the negative distance between TSS and enhancer. The three attention-based scores were evaluated alongside the ABC score and other assays in classifying E-P pairs with significant expression changes, measured using the area under the precision-recall curve (auPRC). To remove gene-related bias, we normalized the scores for all promoter-enhancer pairs, ensuring their sum equals one for each selected gene. These normalized scores were used as the attention scores to assess all enhancer-promoter pairs.

### Motif discovery

We utilized TF-MoDISco^60^ to identify motifs in nucleotide contribution scores across enhancer sequences from the testing set based on the previously determined DeepLIFT contribution scores. We implemented default settings to find seqlet patterns which were then compared against the JASPAR database using Tomtom^61^.To identify motifs of high importance scores, we perform *in silico* saturation mutagenesis (ISM) on the model given a DNA sequence of interest. ISM functions by sequentially substituting each character in a sequence with every other possible character, followed by assessment of the change in the predictive output before and after each substitution. This observed difference is interpreted as a measure of importance or attribution, where a higher magnitude value indicates that the character change has a significant impact on the prediction, thereby suggesting its high importance. Therefore, ISM can be used to uncover block of nucleotides corresponding to TF motifs on the putative enhancer sequence. The corresponding sequence was scanned against the hg38 reference genome, using 2,179 TF motif position weight matrices previously compiled in a global reference map^62^. The cumulative motif weight of the selected motif is calculated by summing the ISM scores of each nucleotide in the fragment. The predicted ISM scores were compared with the corresponding TF ChIP-seq signals derived from ENCODE (Supplementary Table).

## Data availability

All primary experimental datasets used in this study were obtained from publicly accessible sources. The specific repositories and individual accession numbers of these data sources are provided in the Supplementary tables.

## Acknowledgements

This work was supported by National Science and Technology Major Project (No. 2022ZD0115001), National Natural Science Foundation of China (92353304, T2495221) and New Cornerstone Science Foundation (NCI202305). Computations were carried out at Changping Laboratory Supercomputing Center.

